# Targeting the hyperactive STAT3/5 pathway in cutaneous T-cell lymphoma with the multi-kinase nuclear transporter inhibitor IQDMA

**DOI:** 10.1101/2025.03.03.641168

**Authors:** Saptaswa Dey, Helena Sorger, Michaela Schlederer, Isabella Perchthaler, Martin L. Metzelder, Lukas Kenner, Richard Moriggl, Peter Wolf

## Abstract

Cutaneous T-cell lymphoma (CTCL), particularly its tumor stage mycosis fungoides (MF) subtype, presents considerable therapeutic challenges since current treatment modalities show limited efficacy. This study addresses the unmet need for novel targeted therapies that inhibit the STAT3/5 pathway, which is hyperactive in CTCL. Utilizing a murine model with intradermally grafted malignant T-cell lymphoma cells, we compared the efficacy of the multi-kinase inhibitor IQDMA with the conventional, topical psoralen + UV-A (PUVA) phototherapeutic regimen. Our data show that IQDMA reduced tumor volume by 90.7% (*p* = 0.0001) and was significantly more effective than PUVA, which reduced the tumor volume by only 46.2% (*p* = 0.0074). Results of an immunobiological analysis reveal that IQDMA treatment decreased tumor cell infiltration by 29.8% (*p* = 0.03) and the percentage of Ki67^+^ cells by 25.3% (*p* = 0.03), indicating a reduced tumor cell proliferation rate. Moreover, remarkable 40.0% and 45.6% reductions were observed in the total STAT5 (*p* = 0.047) and STAT3 (*p* = 0.01) levels of the infiltrating tumor cells upon IQDMA treatment. STAT5 levels are directly correlated with CD3^+^ tumor cell infiltration, confirming the role of the STAT3/5 pathway in the disease pathogenesis. Intriguingly, while phospho-STAT5 and total STAT5 levels directly correlated in the vehicle-treated group, a negative correlation was observed in the IQDMA-treated group, indicating IQDMA action in blocking STAT5 hyperactivation. IQDMA targets PAK kinase, a nuclear transporter for phospho-STAT5; in turn, we observed a compartmental shift of phospho-STAT5 from the nucleus to the cytoplasm. These key findings establish the properties of IQDMA as a potent targeted therapy for CTCL and offer compelling evidence for its clinical evaluation.

## Introduction

Cutaneous T-cell lymphoma (CTCL) is a heterogeneous group of non-Hodgkin lymphomas characterized by malignant T-lymphocyte proliferation, which primarily affects the skin (1, 2). Delayed or misdiagnosis is common, adversely affecting the disease prognosis. The most common types of CTCL are mycosis fungoides (MF) and Sézary syndrome (SS), although the spectrum also includes less common variants such as primary cutaneous anaplastic large cell lymphoma and lymphomatous papulosis (2, 3). The molecular pathogenesis of CTCL is complex, involving numerous genetic, epigenetic, and microenvironmental factors. (4–7). However, the full spectrum of molecular drivers in CTCL remains incompletely understood, limiting the development of targeted therapies (4, 8–10). Chromosomal abnormalities, such as gains and losses of various chromosomes, have been frequently identified in CTCL patients (4, 11).

Treatment strategies for CTCL depend on the stage of the disease and include skin-directed therapies (topical steroids, phototherapy) (12), systemic therapies (interferon-α, retinoids, methotrexate), and newer targeted therapies (histone deacetylase inhibitors, monoclonal antibodies) (13). In the event of advanced disease states, stem cell transplantation and extracorporeal photopheresis are options (14, 15).

Small molecule inhibitors, multi-kinase inhibitors, antibody-drug conjugates, and other targeted therapies that can specifically address the molecular abnormalities in CTCL cells are in demand (4, 6, 14). These agents could offer more effective and less toxic treatment options than conventional chemotherapy. (14, 16). Personalized medicine in CTCL, however, is still in its infancy. Developing treatment strategies based on individual patient genetics, disease characteristics, and responses to earlier therapies could significantly improve outcomes. This approach requires extensive research involving the genetic profiling and molecular characterization of CTCL (6, 9, 11, 17).

One of the critical molecular pathways deregulated in CTCL is the signal transducer and activator of the transcription (STAT) pathway, which mediates cellular responses to various cytokines and growth factors (18). Among the seven members of the STAT family, STAT3 and STAT5 are the most frequently activated in CTCL (19). STAT3 and STAT5 are involved in regulating various biological processes, such as inflammation, immune response, cell cycle, apoptosis, and angiogenesis (20). The aberrant activation of STAT3 and STAT5 in CTCL can result from mutations, gene amplifications, chromosomal translocations, or dysregulated cytokine signaling (21). Several studies have shown that STAT3 and STAT5 are essential in the pathogenesis and progression of CTCL. For example, STAT3 activation is associated with an increased expression of pro-inflammatory cytokines, such as IL-17 and IL-22, which can promote tumor growth and invasion (22–24). STAT3 activation also confers resistance to apoptosis and enhances the survival of malignant T cells. Moreover, STAT3 activation can modulate the immune microenvironment and suppress the anti-tumor immune response. Similarly, STAT5 activation is implicated in the survival and proliferation of CTCL cells and their resistance to conventional therapies. STAT5 activation can additionally induce the expression of oncogenes (11), such as c-MYC and BCL-XL, and inhibit the expression of tumor suppressors, such as p53 and p21 (25, 26). Furthermore, STAT5 activation can influence T helper cells’ differentiation and function, especially the Th17 subset, contributing to chronic inflammation and tissue damage in CTCL (27, 28). Therefore, STAT3 and STAT5 are key regulators of CTCL pathogenesis and treatment. Their aberrant activation can affect multiple aspects of the disease, such as tumor cell survival, proliferation, differentiation, invasion, angiogenesis, and immune evasion. Thus, targeting STAT3 and STAT5 signaling pathways may offer novel therapeutic strategies for CTCL (4).

Our previous work demonstrated the potential of IQDMA, a novel small molecule multi-kinase inhibitor, to block the STAT3/5 pathway in CTCL cells, suggesting a new avenue for targeted therapy (4). The current manuscript expands on this theme by presenting the results of an evaluation of IQDMA’s efficacy in a C57BL/6 intradermal T-cell lymphoma mouse model. (29) And compare its effects with those of conventional therapies. This approach illustrates IQDMA’s specific targeting capabilities and its superior or complementary role in CTCL treatment.

This study directly addressed a gap in current CTCL treatment by focusing on the mechanistic pathways and evaluating the therapeutic potential of IQDMA. It offers new insights into the role of small-molecule inhibitors in managing CTCL, particularly in stages where conventional therapies are less effective.

## Results

### IQDMA attenuates tumor growth in the C57BL/6 allogenic tumor-stage MF model

To uncover the therapeutic potential and mechanistic insight of the investigational multi-kinase inhibitor IQDMA, we established an allogenic mouse model by intradermally grafting malignant T cells into the backs of C57BL/6 mice and evaluated the therapeutic potential of IQDMA. Mice were shaved on the dorsal surfaces and received intradermal injections of EL4 lymphoma cells to induce tumors; the mice were administered 10 mg/kg of IQDMA or a vehicle control intraperitoneally (IP) daily. In this period, the tumor-emergence,-growth, and survival were monitored along with the body weight of the mice to determine the therapeutic effect of IQDMA (**Figured 1A**). Starting around the fourteenth day, an evident and significant divergence between the two groups was observed, with the IQDMA-treated group exhibiting a marked reduction in tumor growth compared to the control group. This effect was even more pronounced by day twenty: The treated group showed significantly smaller tumor volumes (-49.8%, mean tumor volume 109.2 vs. 54.9 mm^3^), as indicated by the asterisk denoting a *p*-value of 0.01 (**Figure 1B**). Mice in the vehicle (light red line) and IQDMA-treated (light blue line) groups maintained a stable body weight throughout the experiment, indicating that the treatment with IQDMA does not have a significant adverse effect on the overall health of the mice as measured by body weight (**Figure 1C**). In summary, we evaluated the impact of the IQDMA on tumor growth and toxicity in an intradermal allogenic CTCL mouse model.

**Figure 1:**
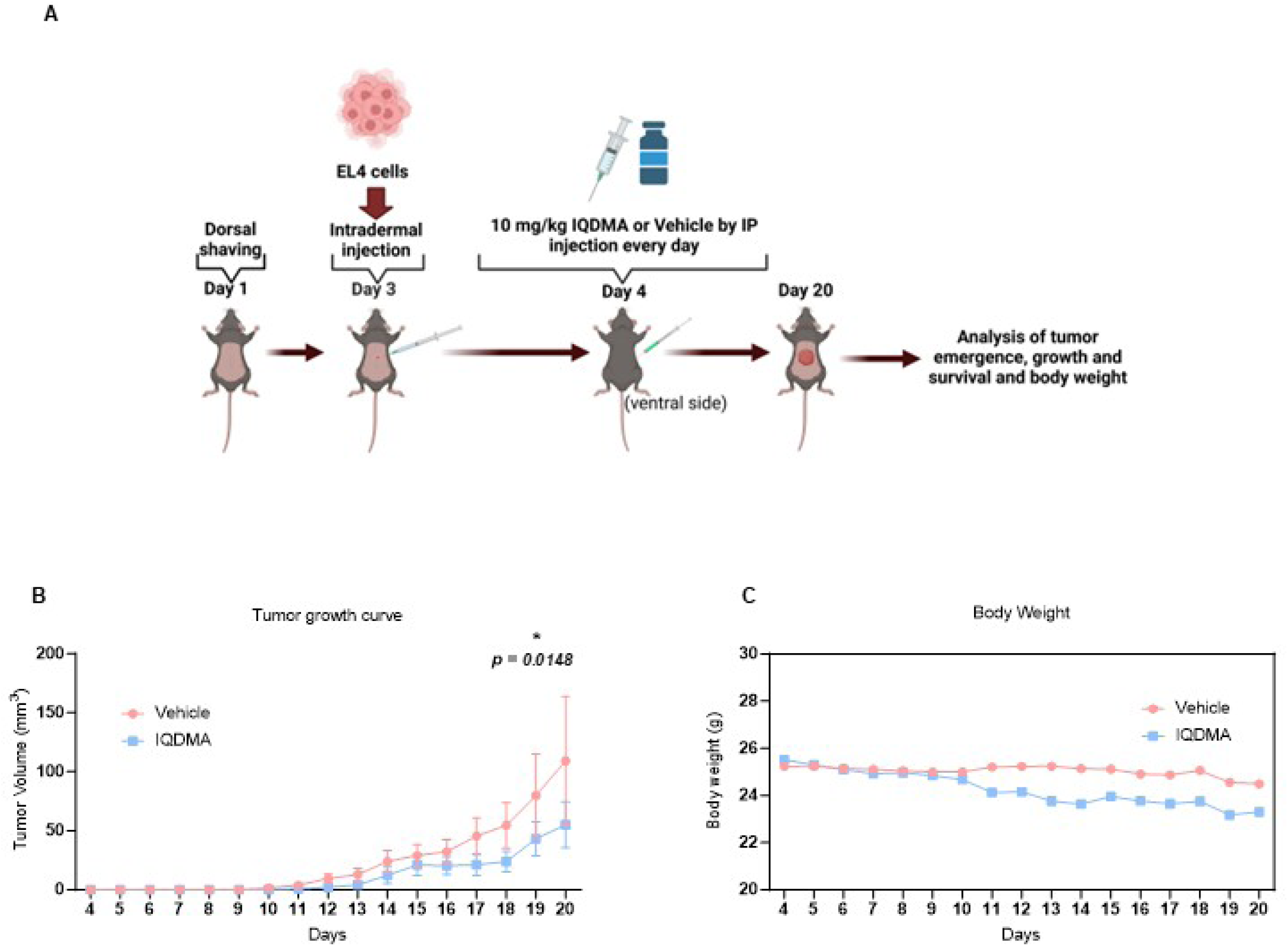
Efficacy of IQDMA in the C57BL/6 intradermal T-cell lymphoma model: **A**: Treatment schema and tumor response in C57BL/6 mice treated with IQDMA. Mice were prepared and injected with EL4 cells, followed by daily IQDMA or vehicle injections. Tumor growth was tracked over time, revealing a lower growth rate in the IQDMA group than in the control, as depicted in the inset plot. **B:** The plot shows tumor volume over time, with the blue line representing the control group and the red line depicting the IQDMA-treated group. Error bars indicate variability within each group, highlighting the therapeutic effect of IQDMA on tumor growth suppression. **C:** Body weight changes in mice throughout the study. The graph compares the average body weight of mice in the control group (blue line) to those in the IQDMA-treated group (red line).

### Assessment of systemic toxicity and histopathological impact of IQDMA treatment in the C57BL/6 model

Mice were administered 10 mg/kg of IQDMA or a vehicle via intraperitoneal injection every day for 14 days to evaluate drug tolerability and toxicity (**Supplementary Figure 1A**). Body weight measurements made at the study’s conclusion revealed no significant difference between the IQDMA and vehicle groups, suggesting that IQDMA does not adversely affect the overall body weight of the mice (**Supplementary Figure 1B**). The blood urea nitrogen levels, a kidney function marker, were comparable between the two groups, indicating no apparent nephrotoxicity due to IQDMA treatment (**Supplementary Figure 1C)**. Aspartate aminotransferase (AST), an enzyme linked to liver health, was similar in IQDMA-treated and vehicle-treated mice, suggesting an absence of hepatotoxicity (**Supplementary Figure 1D)**. Similar alanine aminotransferase (ALT) levels, another enzyme associated with liver function, in both groups further support the absence of liver toxicity due to IQDMA treatment (**Supplementary Figure 1E)**. White blood cell (WBC) count was within normal ranges and comparable between both groups, indicating that IQDMA has no significant immune-toxic effects (**Supplementary Figure 1F)**. There was no significant difference between the two groups of red blood cell (RBC) counts, suggesting that IQDMA does not affect erythropoiesis or cause hemolysis (**Supplementary Figure 1G)**. Hematocrit percentages, a measure of the proportion of blood volume occupied by red blood cells, were also similar between the groups, providing additional evidence that red blood cell mass is unaffected by IQDMA treatment (**Supplementary Figure 1H)**. The hemoglobin levels, essential for oxygen transport, were not significantly different between IQDMA and vehicle groups, suggesting that IQDMA has no impact on the oxygen-carrying capacity (**Supplementary Figure 1I)**. Histological examinations of the kidney, liver, lung, and spleen were performed, and representative micrographs were made for both vehicle and IQDMA-treated groups (Figure 1J). These examinations of the kidney and liver sections revealed no discernible histopathological differences between the two groups and indicated no gross morphological damage. The lung and spleen sections appeared comparable, with no evident histological changes due to IQDMA treatment (**Supplementary Figure 1J)**.

In summary, this evidence indicates that the administration of IQDMA is well-tolerated in healthy C57BL/6 mice, as indicated by the lack of significant changes in body weight and various biochemical markers for organ function. The histological analysis results also reveal no notable pathology in the kidney, liver, lung, or spleen, further supporting the drug’s safety. These findings are critical, as they suggest that IQDMA does not induce systemic toxicity at the dosage evaluated. It might, therefore, be a potential therapeutic agent for treating cutaneous T-cell lymphoma.

### IQDMA outperforms PUVA in the C57BL/6 allogenic intradermal mouse model

Mice underwent dorsal shaving followed by intradermal EL4 lymphoma cell injection. Treatment started when tumors exceeded 1 mm in diameter in at least half of the mice. The experiment had two wings: IP injections of IQDMA or vehicle control and PUVA therapy or no irradiation (**Figure 2A**). Mice that did not receive any irradiation (depicted by the gray line) demonstrated a gradual increase in tumor volume over time (Figure 2). In contrast, the mice that received PUVA therapy (represented by the blue line) exhibited a significant reduction (46.2%) in mean tumor volume (from 126.7 to 68.2 mm^3^, *p-value = 0.0074)* starting on day eight (**Figure 2B**). Mice treated with IQDMA (light blue line) and those treated with the vehicle (light red line) show a striking difference in tumor volume from around day seven, with the IQDMA-treated group showing a significant reduction (90.7%) in mean tumor volume (from 77.6 to 7.2 mm^3^) as compared to the control group (**Figure 2**). This difference becomes highly significant by day ten (*p-value = 0.0001*) (**Figure 2C**).

**Figure 2:**
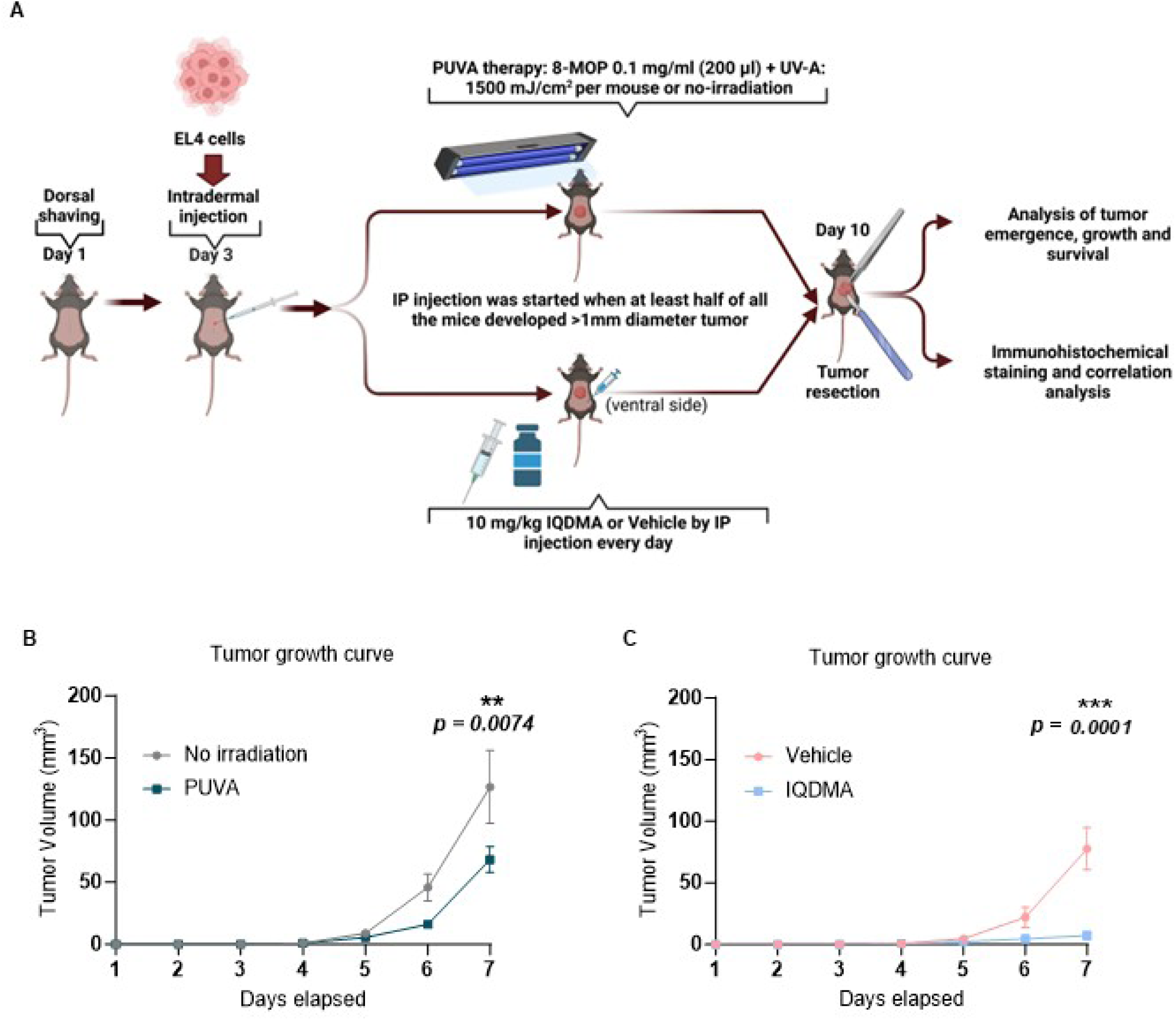
Comparative therapeutic efficacy of PUVA and IQDMA in a T-cell lymphoma model. A: Experimental setup starting with EL4 cell injection, then PUVA therapy or IQDMA treatment, and subsequent tumor resection and analysis. **B:** Tumor volume comparison over time shows PUVA treatment (dark blue line) vs no irradiation (gray line) (*p* = 0.0074). **C:** Tumor volume analysis between IQDMA-treated mice (light red line) and vehicle control (light blue line)(*p* = 0.0001).

Even though both PUVA therapy and IQDMA treatment significantly reduce tumor growth in this mouse model of CTCL, the effect of IQDMA is almost 2 times more pronounced and statistically significant, highlighting its potential as a therapeutic agent.

### A comprehensive analysis of the tumor reveals STAT3/5 dependencies of proliferating tumor cells

Immunohistochemical analysis of the tumor indicates that the tumor cell proliferation correlates directly with their STAT3 and STAT5 positivity status. Hematoxylin and eosin (H&E) staining of tumor sections enabled us to examine the tumor tissue architecture and the infiltration of malignant tumor cells in the intradermal space and measure the perimeter of the infiltrate and its density. (**Figure 3A**). Immunohistochemical staining shows that most of the proliferating malignant cells are CD3^+^ (**Figure 3B**), Ki67^+^ (**Figure 3C**), and STAT3/5^+^ (**Figure 3D, E**). Staining for phosphorylated STAT5 (pY-STAT5) indicates that STAT5 is activated in the proliferating tumor cells (**Figure 3F**), with black arrowheads marking the positive cells).

**Figure 3:**
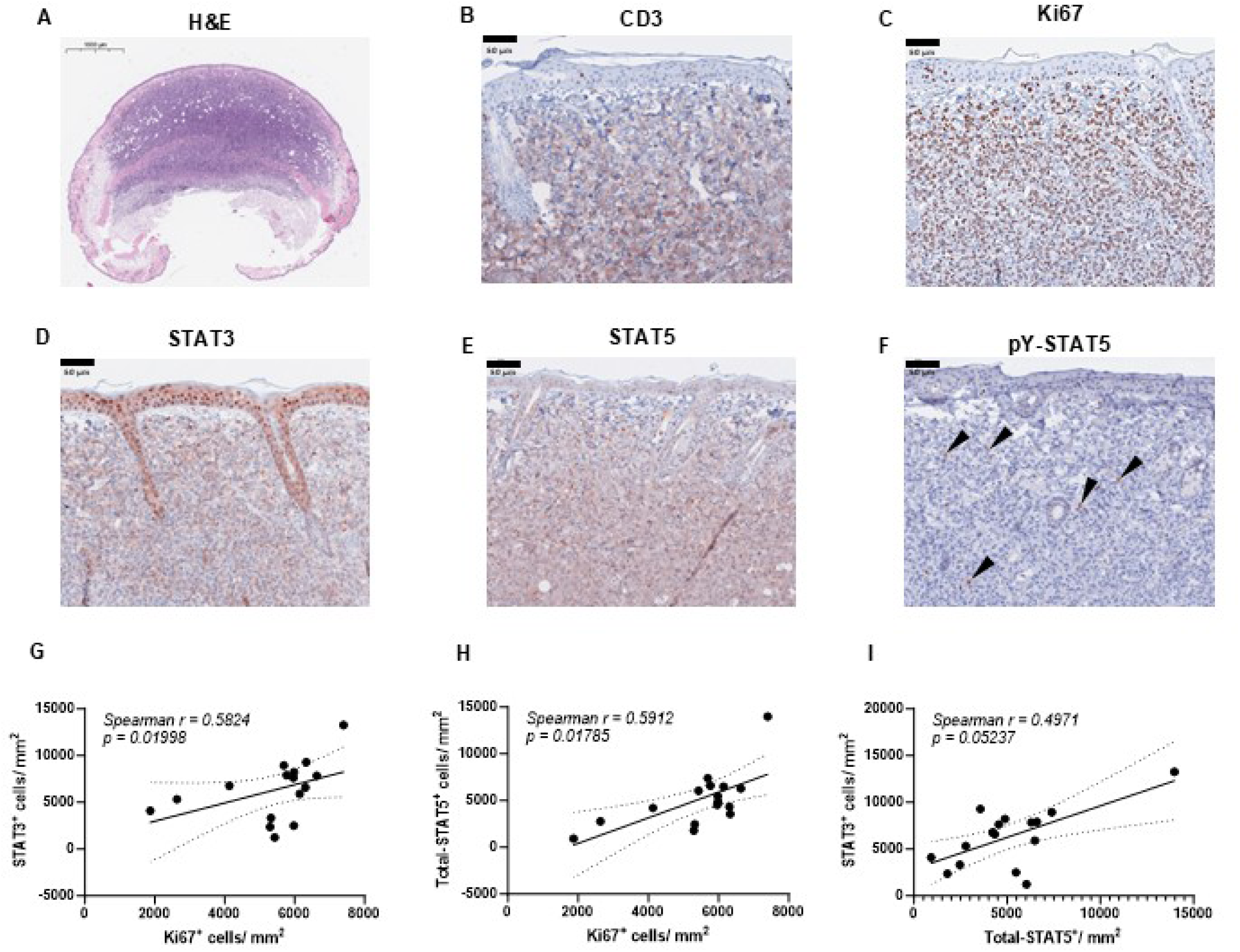
Comprehensive analysis of tumor microenvironment in the T-cell lymphoma model. **A**: H&E staining visualizes the overall tumor architecture. **B**: CD3 staining highlights T-cell infiltration. **C**: Ki67 staining marks proliferative cells. **E-G**: Expression of STAT3, STAT5, and phosphorylated STAT5 (pY-STAT5) (black arrowheads). **G-I**: Correlation analyses between Ki67^+^, STAT3^+,^ and STAT5^+^ cells / mm^2^ (Two-tailed spearmen correlation analysis with 95 % confidence interval, *p* > 0.05).

A significant positive correlation was identified between the number of Ki67^+^ cells and the number of STAT3^+^ cells in tumor tissue, linking STAT3 expression with cell proliferation (**Figure 3G**). A positive correlation was seen between the tumor cell infiltration area and Ki67^+^ cells with total-STAT5^+^ cells, suggesting STAT5 involvement in tumor cell infiltration and cell proliferation (**Figure 3H**). The positive correlation between STAT3^+^ and STAT5^+^ cell numbers (**Figure 3I**) reiterates that the high STAT3/5 expression increases tumor growth in CTCL (4, 30). In summary, our CTCL tumor model provides comprehensive proof of the proliferation and infiltration of malignant T cells in the intradermal space and their direct association with STAT3 and STAT5.

### Kinome-wide inhibition network reveals key hubs targeted by IQDMA

The radial kinome network (**Supplementary Figure 2A**) presents an overview of the inhibitory effects of IQDMA across the entire kinase panel. Nodes represent individual kinases, and edges denote functional and structural similarities, including shared domains and pathway involvement. The color gradient, representing percent control, highlights widespread inhibition, with key hubs such as **ALK, PAK2, AKT1, and STAT3** among the most significantly inhibited. The inhibition of these critical kinases suggests IQDMA’s capacity to target core components of oncogenic signaling networks, particularly within the MAPK, PI3K/AKT, and JAK/STAT pathways. The large number of inhibited kinases within these pathways underscores the multi-target nature of IQDMA and its potential to simultaneously disrupt multiple parallel and redundant signaling networks.

PAK2-centered signaling network highlights its role in linking cytoskeletal dynamics, transcriptional regulation, and survival signaling. The PAK2-specific pathway map (**Supplementary Figure 2B**) demonstrates PAK2 as a central signaling hub with connections to multiple critical pathways. IQDMA-induced inhibition of PAK2 led to predicted downstream effects on its key interactors, including STAT3/STAT5A/STAT5B (nuclear transport and transcription regulation). The interaction between PAK2 and CDC42, along with LIMK1, implicates its involvement in actin cytoskeletal remodeling and cell motility. The inhibition of PAK2 by IQDMA is thus likely to affect multiple cellular processes, including migration, transcriptional regulation, and cell survival.

ALK inhibition disrupts JAK/STAT, MAPK, and PI3K/AKT pathways with downstream implications for proliferation and motility. The ALK-specific pathway map (**Supplementary Figure 2C**) positions ALK as a central regulator of multiple signaling cascades. The inhibition of ALK by IQDMA propagates through downstream pathways, including the JAK/STAT pathway via STAT3, STAT5A, and STAT5B, the MAPK pathway through RAF1 and MAPK1, and the PI3K/AKT pathway via PIK3CA, AKT1, and MTOR. The inhibition of ALK-mediated CDC42 signaling further suggests a potential impact on cytoskeletal organization and cell migration. The extensive downstream network influenced by ALK inhibition highlights its critical role in integrating oncogenic signals, with its inhibition expected to affect cell proliferation, survival, and metastasis.

### IQDMA significantly reduces STAT3 and STAT5 expression in CTCL tumor cells

H&E-stained sections of tumors from the vehicle-treated group and the IQDMA-treated group are shown in (**Figure 4A**, Supplementary Figure 3A) and (**Figure 4B, Supplementary Figure 3B),** respectively. The less-dense tumor cellularity suggests that a reduction in tumor mass occurred following treatment with IQDMA (**Figure 4B**). The difference in tumor cell infiltration between the two groups is quantified in the panel (**Figure 4C**), which shows a significant decrease (29.8%) in the mean infiltration perimeter (from 25,416.6 to 17,847.6 µm) (*p* = 0.03). The reduction in Ki67^+^ cells upon IQDMA treatment compared to the treated group (**Figure 4D and E**) (**Supplementary figure 4A and 4B)** suggests that IQDMA mediated inhibition of tumor cell proliferation. The bar chart (**Figure 4F)** demonstrates a statistically significant reduction of 25.3% (from 6217 to 4642) (*p* = 0.03) of Ki67^+^ cells/mm² in the IQDMA-treated group as compared to the vehicle group.

**Figure 4:**
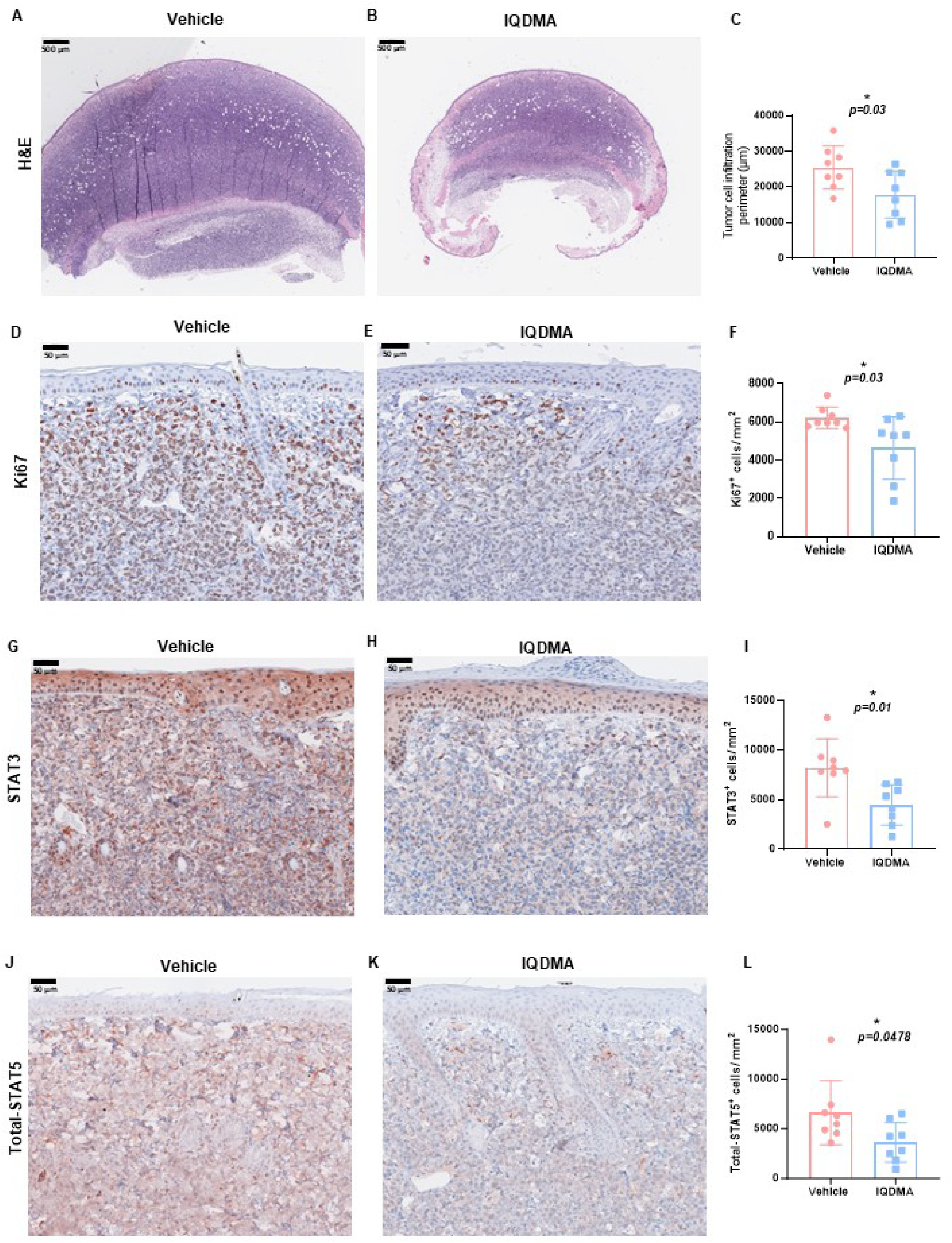
**Immunohistochemical analysis of STAT3 and STAT5 in tumor tissues from the vehicle and IQDMA-treated groups**. **A, B:** A representative image of the vehicle and IQDMA-treated tumors. H&E staining shows differences in tumor cell infiltration and density. **C:** Quantification of tumor cell infiltration area with a significant reduction observed in the IQDMA group. **D, E:** A representative image of Ki67 staining of the vehicle and IQDMA-treated tumors. **F:** Quantification of Ki67^+^ cells. **G, H**: A representative image of STAT3 expression in vehicle and IQDMA treated tumors. **I:** Quantification of STAT3^+^ cells/mm², showing a significant decrease post-IQDMA treatment (*p* = 0.01). **J, K**: A representative image of total STAT5 expression in the vehicle and IQDMA treated tumors. **L:** Quantification of total number of STAT5^+^ cells/mm² indicates a trend towards decreased expression with IQDMA treatment.

Immunohistochemical staining for STAT3 in vehicle-treated tumor tissue (**Figure 4G, Supplementary Figure 5A) and IQDMA-treated tumor tissue (Figure 4H, Supplementary Figure 5B**) revealed a marked decrease in STAT3 expression upon IQDMA treatment compared to the vehicle group. A 45.6% reduction of STAT3^+^ cells/mm^2^ (from 8199 to 4457) (*p* = 0.01) (**Figure 4I**).

Furthermore, immunohistochemical staining for total-STAT5 in vehicle-treated (**Figure 4J, Supplementary Figure 6A) and IQDMA-treated (Figure 4K, Supplementary Figure 6B**) tumor tissues depicted a visible reduction in STAT5 expression upon IQDMA treatment. A bar chart shows a decrease of 40.0% in the average number of STAT5^+^ cells/mm^2^ (from 6590 to 3630) (*p* = 0.0478) in the IQDMA-treated group as compared to the vehicle group (**Figure 4L**).

The decrease in the expression of STAT3 and STAT5 measured in the tumor tissues following IQDMA treatment suggests that the drug exerts its therapeutic effects by modulating these key signaling pathways involved in cell proliferation and survival. The results shown in Figure 4 support the potential of IQDMA as a targeted therapeutic agent in CTCL due to its pronounced effects on the STAT signaling pathways in the tumor tissue.

### Subcellular localization shifts in pY-STAT5 signal the effects of IQDMA treatment

Representative images of immunohistochemical staining for cytoplasmic and nuclear pY-STAT5 in the vehicle-treated and IQDMA-treated tumor cells are shown in **Figure 5A, Supplementary 7 Figure 7A** and **Figure 5B, Supplementary Figure 7B**, respectively, with the arrowheads marking positive staining. A comparison of panel A vs. B demonstrates a decrease in nuclear pY-STAT5 localization in the IQDMA-treated cells, reinforcing the suggestion that IQDMA could inhibit STAT5-mediated transcriptional activation by retaining it in the cytoplasm. (4)These IHC results indicate that IQDMA treatment affects the localization of pY-STAT5 in the tumor cells by increasing the cytoplasmic retention of pY-STAT5, as noted in the staining intensities, and decreasing the nuclear presence of pY-STAT5 (**Figure 5A-B and Supplementary Figure 6A-B**). This suggests a diminished role of STAT5 in the nucleus, possibly leading to reduced transcriptional activity of the STAT5 target genes. (4), leading to reduced tumor cell proliferation and increased survival.

**Figure 5:**
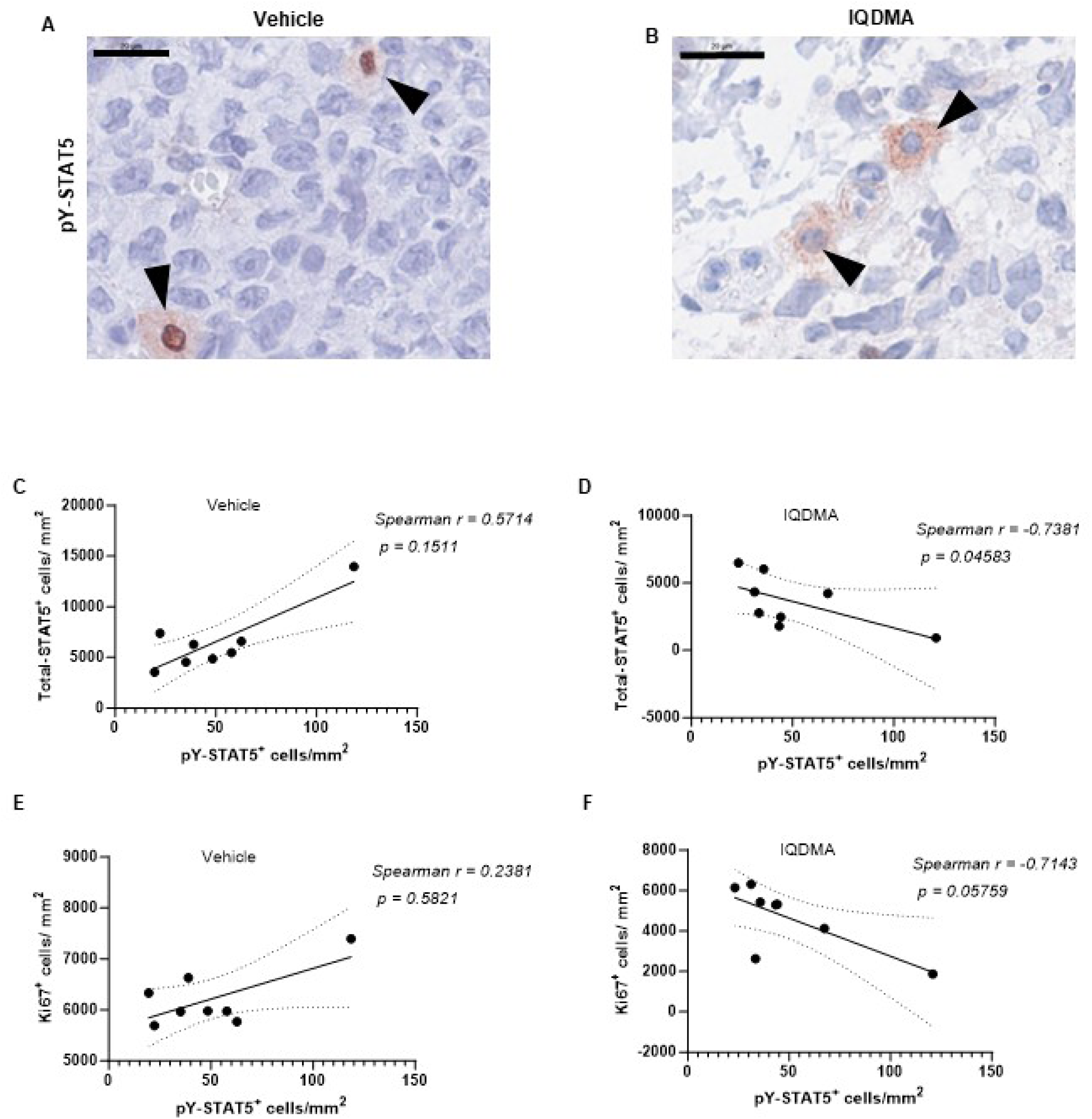
Localization and quantification of phosphorylated STAT5 (pY-STAT5) in vehicle vs. IQDMA-treated cells. A-B: Immunohistochemical staining of pY-STAT5, showing nuclear and cytoplasmic (**A**, black arrowheads) and only cytoplasmic localization (**B**, black arrowheads). **C-D:** Correlation between total number of STAT5^+^ and pY-STAT5^+^ cells/mm², with the vehicle group showing positive correlation (*r* = +0.57, *p* = 0.07) and IQDMA group showing negative correlation (*r* =-0.66, *p* = 0.04). **E-F:** The negative correlation between the number of Ki67+ cells/mm² and pY-STAT5+ cells/mm² is more pronounced in the IQDMA group (*r* =-0.71, *p* = 0.02), (Two-tailed spearmen correlation analysis with 95 % confidence interval, *p* > 0.05).

These observations collectively suggest that IQDMA exerts its anti-tumor effects by modifying the intracellular distribution of pY-STAT5, preventing its nuclear localization, which is critical for cancer cell growth and survival.

### IQDMA disrupts STAT5 activation and tumor proliferation while modulating immune infiltration in CTCL

We explored the relationships between pY-STAT5 levels and various tumor parameters in our mouse model of cutaneous T-cell lymphoma, comparing the vehicle vs. IQDMA treatment. A moderate positive correlation (*r* = +0.57) is seen between the total STAT5 and pY-STAT5 levels in the vehicle-treated group (**Figure 5C**), suggesting that a higher total STAT5 level is associated with higher levels of its active form, pY-STAT5. Conversely, these levels in the IQDMA-treated group (**Figure 5D**) are significantly negatively correlated (*r* =-0.73, *p* = 0.04), indicating that IQDMA treatment disrupts the relationship between the total and phosphorylated STAT5 levels, potentially affecting STAT5’s role in tumor progression.

The correlation between the cell proliferation, as indicated by the Ki67^+^ cell, and pY-STAT5 levels in the vehicle group (**Figure 5E**) is weak and positive (*r* = +0.23). In contrast, these levels display a strong negative correlation (*r* =-0.71, *p* = 0.05) in the IQDMA group (**Figure 5F**), suggesting that IQDMA treatment inhibits tumor cell proliferation by affecting STAT5 phosphorylation.

Taken together, these findings indicate that IQDMA’s role in disrupting the nuclear transportation of pY-STAT5, thereby hindering the pathogenic STAT3/5 pathway, is key for the proliferation and survival of CTCL cells.

## Discussion

The quest to elucidate effective therapeutic strategies for cutaneous T-cell lymphoma (CTCL) is a journey fraught with complexities and challenges. This study examined the potential of IQDMA as a targeted therapeutic agent for disrupting the STAT3/5 pathway. Still, our narrative remains incomplete without acknowledging the inherent limitations and anticipating the future trajectory of our scientific endeavor.

IQDMA targets key signaling hubs, which has implications for network-based therapeutic strategies. The kinome-wide network analysis demonstrates that IQDMA effectively targets critical signaling hubs, including ALK, PAK2, and STAT3/5, all of which are known to play pivotal roles in oncogenesis (31, 32). The inhibition of these central nodes disrupts multiple parallel pathways, reducing the likelihood of single-pathway compensation and potentially minimizing resistance development. ALK and PAK2, in particular, emerge as key regulators within the network, each influencing several downstream signaling components. ALK’s involvement in JAK/STAT, MAPK, and PI3K/AKT pathways places it at the intersection of cell growth, survival, and motility, while PAK2’s connections to cytoskeletal dynamics, nuclear transportation, and transcriptional regulation make it a key driver of both structural and gene-expression-related processes (33, 34).

While the inhibition of central nodes such as ALK and PAK2 suggests a broad impact on signaling networks, it also raises the possibility of feedback loops and compensatory pathway activation (24, 35, 36). For example, inhibition of PAK2 and its downstream effectors may trigger compensatory activation of parallel pathways, such as PI3K/AKT or Rho-family GTPases, potentially restoring some level of cell motility or survival (33, 37, 38). Similarly, ALK inhibition may result in signal redirection through alternative receptor tyrosine kinases (RTKs) or downstream effectors (8, 39, 40). These compensatory mechanisms highlight the need for combination therapies, where IQDMA could be paired with inhibitors targeting complementary pathways or feedback mechanisms to achieve a more durable therapeutic response.

The network-based targeting by IQDMA provides a strong rationale for clinical use of this drug. Simultaneous inhibition of ALK and PAK2 blocks the hyperactivation of STAT3, STAT5A, and STAT5B and the downstream regulators of STAT pathways such as MYC. PIM1, MCL1, Cyclin D2, Cyclin D3, etc., address the redundancy and resistance mechanism often observed in oncogenic signaling networks (36, 41, 42). By disrupting multiple key pathways simultaneously, IQDMA may reduce the likelihood of adaptive resistance. However, identifying potential secondary targets—such as kinases in feedback loops or alternative pathways—is essential to optimize therapeutic efficacy. Given its broad impact, IQDMA could be combined with inhibitors of parallel or compensatory pathways, such as SRC or Rho-family kinases, to achieve synergistic effects (43).

Dynamic simulations of signaling propagation in the context of kinase inhibition could help identify key nodes likely to mediate resistance (44). In future work, combining computational models with experimental phosphoproteomic data could refine predictions and prioritize targets for combination therapies. Additionally, further investigation of the impact of IQDMA on tumor cells with different genetic backgrounds will be essential to determine its effectiveness across various cancer types and to identify biomarkers of response or resistance (45–47).

The intrinsic limitation of animal models in recapitulating the human condition. Despite its utility, the biological milieu of a murine system only partially mirrors the intricate immunological and genetic landscape of human CTCL. This discrepancy raises questions about the direct applicability of our findings to the clinical setting, as the interplay between tumor and host in humans is a tapestry woven with far more immune complexity.

Furthermore, while shedding light on the role of STAT signaling, our study’s focus on a single pathway might obscure other contributing molecular pathways like the gain of MYC locus or loss of p53 locus (4). CTC L’s intricacy, characterized by its heterogeneity and multifaceted nature, means that our understanding remains superficial without considering the broader network of cellular signals and tumor microenvironment interactions (48).

Targeted therapies such as mogamulizumab (anti-CCR4) (49), alemtuzumab (anti-cd52) (50–52), and brentuximab-vedotin (Anti-CD30) (53–55) have revolutionized the treatment of CTCL, offering more efficacy and tolerability than conventional therapies. However, continued research and innovation are crucial to addressing unmet needs, such as drug tolerability, preventing disease relapse, and improving patients’ lives with CTCL. CTCL, mainly focusing on its challenges and the future implications of targeted agents for STAT3 and STAT5 proteins, can be informed by several research papers (4, 8, 56–58).

CTCL is a complex disease, and its pathogenesis remains unclear. However, recent research has shed light on potential therapeutic targets, including the STAT3 and STAT5 proteins. CTCL patients show elevated levels of the oncogenic microRNA miR-155 (59). STAT5, which is abnormally activated in malignant T cells, drives the expression of miR-155. This suggests that the STAT5/miR-155 pathway promotes the proliferation of malignant T cells and could be a target for therapy in CTCL (19). In human CTCL cell lines, specifically, the knockdown of STAT3 induces apoptosis and downregulates anti-apoptotic genes like Bcl-xL. This indicates that STAT3 also is crucial for the survival of CTCL cells and may serve as a novel therapeutic target (60). Furthermore, both STAT3 and STAT5 are known to contribute to drug resistance in hematopoietic malignancies. The development of inhibitors targeting these proteins has been the subject of intense investigations, suggesting their potential in CTCL therapy (58). In leukemic CTCL, copy number gains of STAT3/5 correlate with increased clonal T-cell count. Inhibiting these pathways could be an effective therapeutic strategy (4).

The lack of small molecule agents for CTCL is partly due to the inadequate understanding of the CTCL transcriptional and translational landscape and the scarcity of specific molecular agents (56). IQDMA, a multi-kinase small molecule inhibitor, affects tumor growth, cell proliferation, apoptosis, and STAT signaling in the mouse model of this study. Our data demonstrates that IQDMA treatment significantly reduces tumor volume. Our key findings include a marked decrease in tumor cell proliferation, as indicated by reduced Ki67 expression, upon IQDMA treatment compared to the vehicle. A reduction in the expression of STAT3 and STAT5 suggests that IQDMA disrupts these critical signaling pathways. Finally, significant changes in the subcellular localization of pY-STAT5, with increased cytoplasmic retention and decreased nuclear presence in IQDMA-treated tumors, reflect its ability to disrupt the nuclear transportation of STAT3/5.

These findings suggest that IQDMA has a significant therapeutic effect on CTCL by inhibiting tumor cell proliferation and modulating STAT3/5 signaling pathways by disrupting their nuclear transportation.

## Materials and Methods

### Animal Model and Housing Conditions

C57BL/6 mice (strain Ncrl, 4–6 weeks old) were sourced from Charles River Laboratories, Germany (Germany GmbH). The mice were housed under pathogen-free barrier conditions in individually ventilated cages, maintained on a 12-hour light/dark cycle with access to standard food and water ad libitum. Animal handling and experimentation were performed in compliance with institutional guidelines on animal welfare, with ethical approval granted by the Federal Ministry of Science, Research, and Economy of Austria (approval number: BMBWF-66.010/0064-V/3b/2018). Mice were randomly assigned to treatment groups based on sex and age, with a minimum of 8 animals per group to ensure adequate statistical power.

### Tumor Cell Preparation and Injection

EL4 murine T-cell lymphoma cells were cultured under standard conditions, and an appropriate cell suspension was prepared for intradermal injection. On Day 1, the dorsal region of each mouse was shaved to allow direct access for tumor cell grafting. EL4 cells were intradermally injected into the shaved area on Day 3, with tumor initiation monitored daily. Tumor development was observed, and subsequent treatments were initiated once tumor size exceeded 1 mm in diameter in at least 50% of the mice in each group.

## Photochemotherapy (PUVA) Treatment

For the PUVA-treated groups, photochemotherapy was initiated using 8-methoxy psoralen (8-MOP) in combination with UVA radiation. Mice received 200 μL of 8-MOP at a 0.1 mg/mL concentration via intraperitoneal (IP) injection. Subsequently, animals were exposed to UVA light at an intensity of 1500 mJ/cm² per mouse using a UVA irradiation device (details regarding device model and specific wavelength, if applicable, should be inserted here). Mice in the control group were injected with 8-MOP but were not exposed to UVA irradiation. This protocol was conducted to evaluate the impact of photochemotherapy on tumor growth and emergence in the EL4 model.

### Pharmacologic Intervention with IQDMA

Mice in the experimental group were administered IQDMA daily at a dose of 10 mg/kg dissolved in a vehicle solution comprising 5% dimethyl sulfoxide (DMSO), 50% polyethylene glycol (PEG-400), 5% TWEEN®80, and 40% distilled water. IP injections were consistently administered to both the treatment and vehicle control groups from Day 4 post-tumor injection. This pharmacologic intervention was maintained until Day 20, when tumor growth and treatment effects were assessed through volumetric analysis and histological evaluation.

### Tumor Growth Assessment

Tumor growth was assessed daily from Day 4 post-injection until Day 20. Tumor dimensions (length and width) were measured with Vernier calipers, and tumor volumes were calculated using the formula:

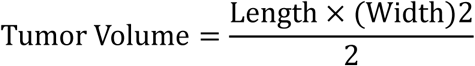

### Body Weight Monitoring

Body weight was recorded daily throughout the experimental period to indicate general health and treatment tolerability. Any significant weight fluctuations were noted, particularly to assess the potential toxicity of the IQDMA treatment compared to the vehicle control.

### Drug Tolerability and Toxicity Studies

An independent cohort of C57BL/6 mice was used to evaluate the tolerability and potential toxicity of IQDMA. Animals were maintained under pathogen-free housing conditions as previously described. Mice in the experimental group received daily intraperitoneal (IP) injections of IQDMA at 10 mg/kg, consistent with previous treatment regimens, to evaluate drug tolerability over 14 days. Body weight was recorded daily as a general indicator of health and drug tolerability. Additionally, all mice were closely monitored for any overt clinical signs of distress or toxicity, including lethargy, changes in coat condition, and signs of inflammation or infection.

### Blood Biochemistry and Hematology Analysis

On Day 14, blood samples were collected from mice for hematological and biochemical analyses to assess the impact of IQDMA on systemic physiology. Key parameters were measured, including blood urea nitrogen (BUN), aspartate aminotransferase (AST), alanine aminotransferase (ALT), white blood cell (WBC) count, red blood cell (RBC) count, hematocrit (HCT), and hemoglobin (Hb) levels. These parameters were chosen to monitor potential effects on renal and hepatic function and general hematologic health.

### Histopathological Examination

Following blood collection on Day 14, mice were euthanized, and tissues, including the kidney, liver, lung, and spleen, were harvested for histopathological evaluation. Tissues were fixed in 10% neutral-buffered formalin, processed, and embedded in paraffin. Sections (4 μm thick) were stained with hematoxylin and eosin (H&E) for morphological assessment. Microscopic examination was conducted to identify any potential tissue-specific toxicities, focusing on inflammation, cellular damage, and architectural changes in the organ tissues of IQDMA-treated mice compared to vehicle controls.

### Tumor Resection and Histological Analysis

Tumors reaching the 10 mm size threshold were resected for histopathological evaluation. Tissue sections were prepared and stained for immunohistochemical analysis to correlate treatment-induced changes in tumor morphology and immune cell infiltration. Details regarding specific antibodies, staining procedures, and imaging techniques used for analysis should be documented in the subsequent section as per laboratory protocol.

### Immunohistochemical Analysis of Tumor Biopsies

To investigate the cellular and molecular characteristics within tumor biopsies, immunohistochemical (IHC) studies were conducted on 2-μm-thick sections of formalin-fixed, paraffin-embedded (FFPE) skin tissue. Following sectioning, slides were deparaffinized in xylene and rehydrated through graded ethanol solutions to prepare for antigen retrieval and antibody staining.

### Antigen Retrieval and Immunostaining Protocol

Antigen retrieval was performed by incubating slides in TRIS-EDTA buffer (pH 9.0) to unmask epitopes and enhance antibody binding. After cooling, sections were blocked and subsequently incubated with primary antibodies specific to target proteins at optimized concentrations. The following primary antibodies were used: **CD3** (Rabbit monoclonal; #RM9107-S0, Thermo Fisher Scientific, USA), a marker for T-cell infiltration, **STAT3** (Clone 9D8; #MA1-13042, Thermo Fisher Scientific), which plays a role in tumor-associated inflammation and proliferation, **STAT5A** (Clone E289; #ab32043, Abcam, UK) and **STAT5B** (#ab235934, Abcam), markers involved in JAK-STAT signaling associated with immune response and cell survival, **pY-STAT5** (Phospho-Tyr694/699; #9359, Cell Signaling Technology, USA), a phosphorylated form of STAT5 indicative of activation.

The slides were incubated with the primary antibodies according to the manufacturer’s instructions, followed by detection using an appropriate secondary antibody conjugated to a chromogenic substrate.

### Imaging and Quantitative Analysis

Stained slides were digitized using the Panoramic Digital Slide Scanner (3DHistech Ltd., Hungary) and the Aperio Digital Pathology Slide Scanner (Leica Biosystems, Germany) to capture high-resolution images of the tissue sections. Digital IHC images were subsequently analyzed using CaseViewer software (3DHistech Ltd.) and QuPath (v0.2.3), an open-source platform designed to analyze whole slide images quantitatively. Parameters such as CD3+ T-cell density, Ki67 proliferation index, and STAT3/Total-STAT5 expression were quantified in representative tumor regions, and associations were analyzed statistically. Scripts are deposited to the following GitHub repository: https://github.com/SaptaDey/QuPath-Posetive-cell-detection/tree/main.

### Kinome Data Preprocessing

Kinase inhibition data was extracted and analyzed from our previously published kinome-screen dataset in our EMBO MolMed manuscript: supplementary meterials: dataset EV4: https://doi.org/10.15252/emmm.202115200. The screen was performed using a panel of kinases, and percent control values were calculated for each kinase, representing residual activity compared to untreated controls. Data preprocessing involved filtering non-redundant kinases and standardizing percent control values on a scale from 0 (complete inhibition) to 100 (no inhibition). Missing or incomplete data were excluded prior to downstream analysis.

### Network Mapping

The kinome network was constructed to visualize the overall inhibition profile across the entire kinase panel following IQDMA treatment. The network was generated using NetworkX in Python, with nodes representing individual kinases and edges denoting functional or structural similarities, such as kinase family relationships or shared catalytic domains, curated from kinase classification resources. The network layout followed a circular (radial) topology to facilitate visual interpretation. Node attributes, including size and color, were mapped to percent control values to highlight key kinases and their inhibition status. Highly inhibited kinases (low percent control) were visually emphasized using a continuous color gradient (Viridis colormap) ranging from blue (strong inhibition) to yellow (minimal inhibition). The network was plotted using Matplotlib, and the final figure was exported at 300 dpi for publication.

### Pathway Mapping

To explore key IQDMA targets within specific signaling contexts, pathway-specific maps were generated for PAK2 and ALK using kinase inhibition data and curated protein–protein interaction networks from KEGG, Reactome, and STRING databases. Interactions between PAK2 and transcriptional regulators STAT3 and STAT5A/STAT5B were mapped to emphasize its role in nuclear transport and transcriptional activation. The ALK-centered network included its downstream interactions with the MAPK (RAF1, MAPK1), PI3K/AKT/mTOR (PIK3CA, AKT1, MTOR), and JAK/STAT (STAT3, STAT5A, STAT5B) pathways. The network was constructed using NetworkX in Python, with nodes and edges representing kinases and their functional connections. Node size and color were mapped to percent control values using the same colormap as **Supplementary Figure 2A**. Both pathway-specific maps were visualized using Matplotlib, and final figures were exported at 300 dpi.

### Dynamic Signal Flow and Network Propagation Analysis

Dynamic pathway behavior and downstream effects of kinase inhibition were modeled using a system of ordinary differential equations (ODEs). For each pathway, differential equations were used to simulate signal propagation and feedback mechanisms. The rate of inhibition decay and activation of downstream effectors was modeled using Python’s **SciPy** library, specifically the odeint function for numerical solutions. Dynamic signal propagation was used to identify key nodes (hubs) and bottlenecks within the network.

### Software and Tools

- **Python (v3.9)**: Used for network construction, data processing, and visualizations
- **NetworkX**: Used for creating node-edge diagrams and mapping kinase interactions
- **Matplotlib**: Used for static network visualizations and figure generation
- **SciPy**: Used for ODE-based signal propagation modeling
- **Pandas**: Used for data manipulation and preprocessing
- **KEGG, Reactome, and STRING**: Used for pathway and functional annotation of kinases
- **R studio 12.0** and **R 4.4.2** (https://cloud.r-project.org/)

All figures were generated in Jupyter Notebooks and exported as high-resolution images (300 dpi) using **Matplotlib’s** savefig() function. The computational analysis pipeline was designed to be fully reproducible and scalable to larger kinase datasets if necessary.

## Statistical Analysis

Data was analyzed using a statistical software package (e.g., GraphPad Prism, python or R). Tumor growth curves were analyzed using an unpaired t-test or ANOVA, as appropriate. P-values were adjusted for multiple comparisons, with a significance threshold at p < 0.05. Data from IHC analyses were subjected to two-tailed Spearman correlation analysis to evaluate associations between cell densities (e.g., CD3+, Ki67+ cells) and molecular markers (e.g., STAT3, STAT5, and phosphorylated STAT5). Spearman’s correlation tests were applied, and significance levels were set at p < 0.05.

## Finding

Austrian Science Fund (FWF - W1241): “DK-MOLIN” - (Molecular Basis of Inflammation) program, which is funded in equal parts by the Doctoral Program funding line of the FWF – (W1241) and Medical University of Graz (SD).

Austrian Science Fund grant (FWF) SFB F04707-B20 (HS, RM).

## Author contributions

Conceptualization: SD

Resources: PW

Data curation: SD

Software: SD

Formal Analysis: SD, HS

Supervision: PW

Funding acquisition: PW

Validation: SD

Investigation: SD

Visualization: SD and HS

Methodology: SD, HS, MS, IP.

Writing – original draft: SD and PW

Project administration: PW

Writing – review & editing: SD, HS and PW.

## Acknowledgments

We wish to thank the Department of Biomedical Research (BMF) at the Medical University of Graz and the Unit of Functional Cancer Genomics at the University of Veterinary Medicine Vienna (Vienna, Austria) for Providing animal husbandry and helping to organize mouse experiments. We would also like to thank Andrea Teufelberger and Theresa Benezeder for their guidance and scientific discussions while formulating the manuscript. We used large language models (LLM) to assist in drafting portions of the introduction and discussion sections, refining language, and checking grammar. The authors reviewed and edited the LLM-generated content for accuracy and coherence.

## Supplementary Material

**Supplementary Figure 1:**
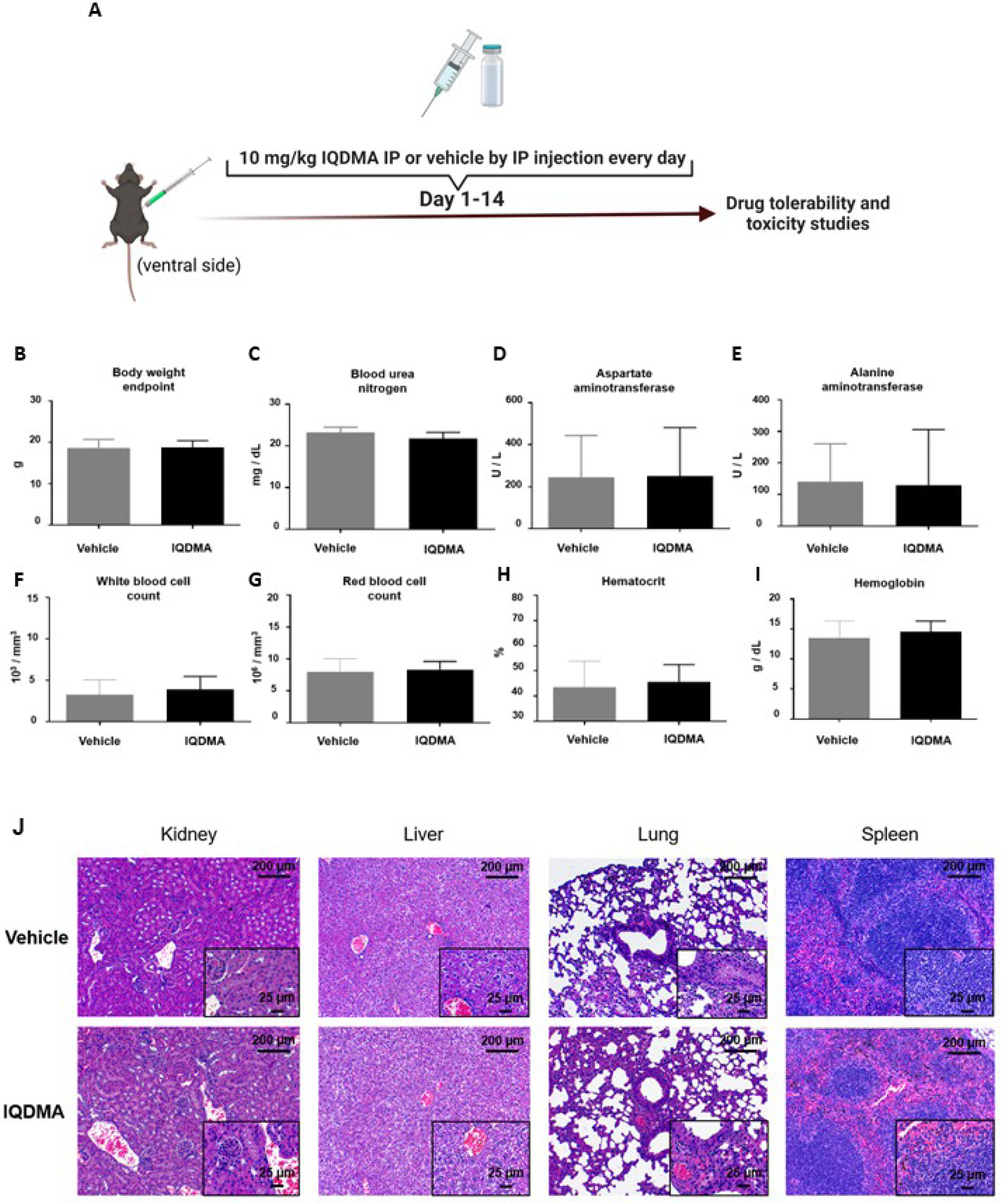
Evaluation of tolerability and toxicity of IQDMA in mice following intraperitoneal administration. **(A)** Schematic representation of experimental design. Drug tolerability and toxicity were assessed at the end of the treatment period. **(B-I)** Assessment of various physiological and hematological parameters at the endpoint: **(B)** Body weight (g), **(C)** Blood urea nitrogen (mg/dL), **(D)** Aspartate aminotransferase (U/L), **(E)** Alanine aminotransferase (U/L), **(F)** White blood cell count (10⁶/mm³), **(G)** Red blood cell count (10⁶/mm³), **(H)** Hematocrit (%), and **(I)** Hemoglobin (g/dL). Data are presented as mean ± standard deviation (SD), with no significant differences observed between the vehicle and IQDMA-treated groups. **(J)** Histopathological analysis of vital organs, including the kidney, liver, lung, and spleen, following hematoxylin and eosin (H&E) staining. Representative images are shown for both vehicle-and IQDMA-treated mice at 200 μm and 25 μm magnifications (insets). No apparent pathological changes were observed between the groups, indicating the absence of significant tissue toxicity.

**Supplementary Figure 2:**
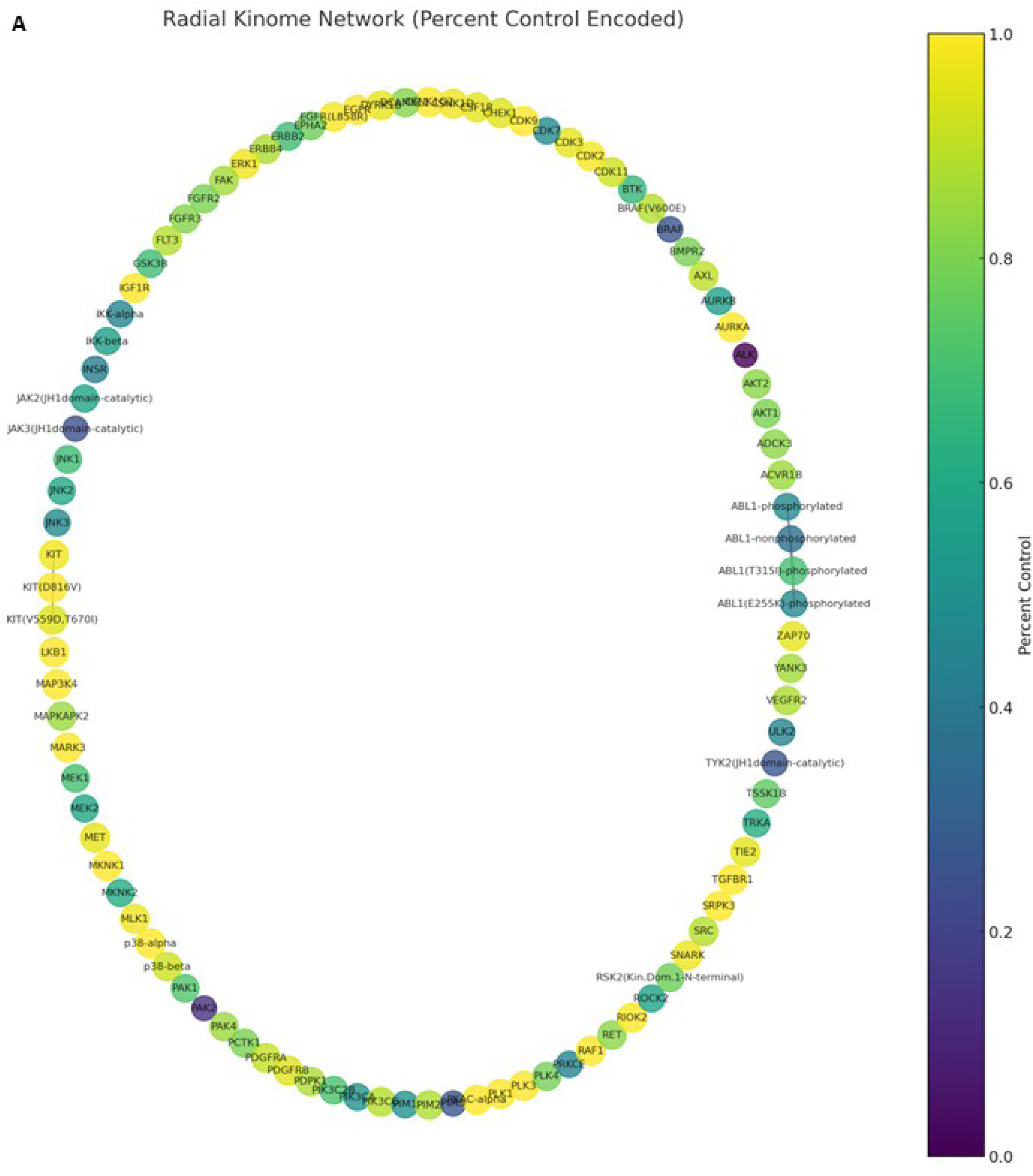
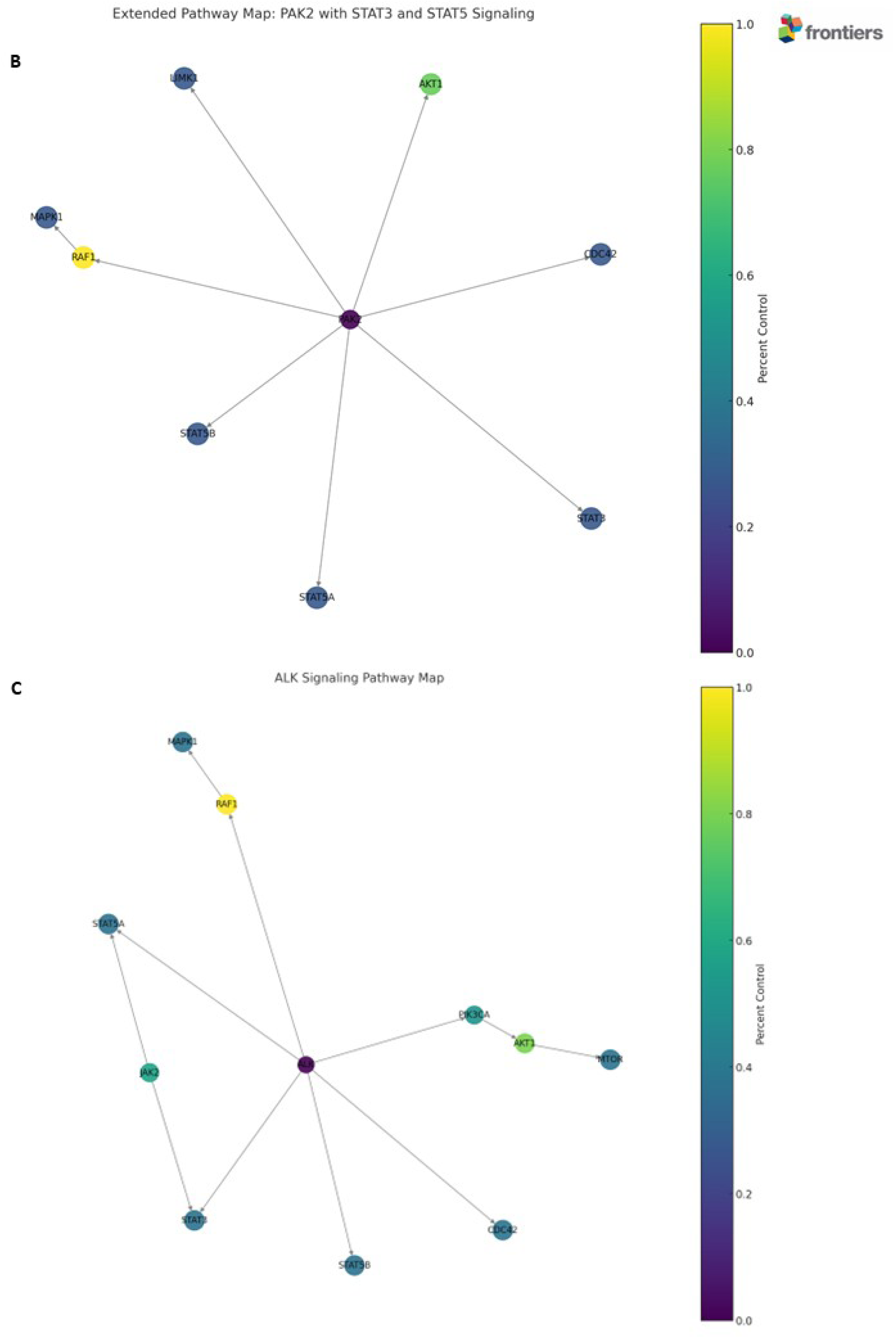
Kinome network and pathway-specific maps illustrating the effect of the multi-kinase inhibitor IQDMA on key signaling networks. **(A)** This radial network visualization illustrates the percent inhibition (percent control) of kinases following treatment with the investigational small-molecule multi-kinase inhibitor IQDMA (Compound Concentration 10 µM). Each node in the network represents an individual kinase, with edges connecting kinases based on functional or structural similarity. The size and position of nodes within the circular layout do not encode specific quantitative information but facilitate visualization of the overall network. The color gradient (from blue to yellow) corresponds to the percent control of kinase activity, as indicated by the color bar. Blue nodes represent highly inhibited kinases (low percent control), whereas yellow nodes indicate kinases with minimal inhibition. Notable hubs such as ALK, AKT1, PAK2, and key kinases involved in pathways like MAPK, JAK/STAT, and PI3K/AKT, are prominently featured, highlighting potential primary and off-target interactions. This figure underscores the broad-spectrum impact of IQDMA and supports its potential for network-based therapeutic applications. **(B)** Extended pathway map illustrating the role of PAK2 and its downstream connections to STAT3 and STAT5 signaling following treatment with the multi-kinase inhibitor IQDMA. Nodes represent individual kinases, and node colors indicate percent control (inhibition) levels, as shown by the color bar. Blue to green nodes represent higher inhibition, while yellow nodes indicate lower inhibition. PAK2 is positioned as a central hub interacting with key signaling pathways: (1) nuclear transport of STAT3, STAT5A, and STAT5B, supporting transcriptional regulation, (2) interaction with RAF1 and MAPK1 in the MAPK/ERK pathway, linking cytoskeletal regulation to proliferation, (3) interaction with AKT1 in the PI3K/AKT pathway, contributing to cell survival, and (4) interaction with CDC42 and LIMK1, emphasizing its role in actin remodeling and cell migration. This diagram highlights the potential of IQDMA to modulate multiple interconnected pathways through PAK2, impacting both upstream and downstream effectors. (**C)** ALK signaling pathway map depicting downstream interactions and pathway modulation following treatment with the multi-kinase inhibitor IQDMA. Nodes represent individual kinases, and node colors reflect the degree of inhibition (percent control) as indicated by the color bar, with blue to green nodes showing higher inhibition and yellow nodes indicating lower inhibition. ALK is positioned as a central hub interacting with multiple key signaling pathways, including: (1) MAPK/ERK signaling via RAF1 and MAPK1, promoting cell proliferation and differentiation, (2) PI3K/AKT/mTOR pathway via PIK3CA, AKT1, and MTOR, regulating cell survival and metabolism, (3) JAK/STAT pathway through direct activation of STAT3, STAT5A, and STAT5B, modulating transcriptional programs, and (4) cytoskeletal regulation through interactions with CDC42, impacting cell migration and morphology. These pathway maps collectively demonstrate the broad impact of IQDMA on key kinases and signaling networks, emphasizing its potential for disrupting oncogenic pathways and suggesting candidate nodes for combination therapies.

**Supplementary Figure 3:**
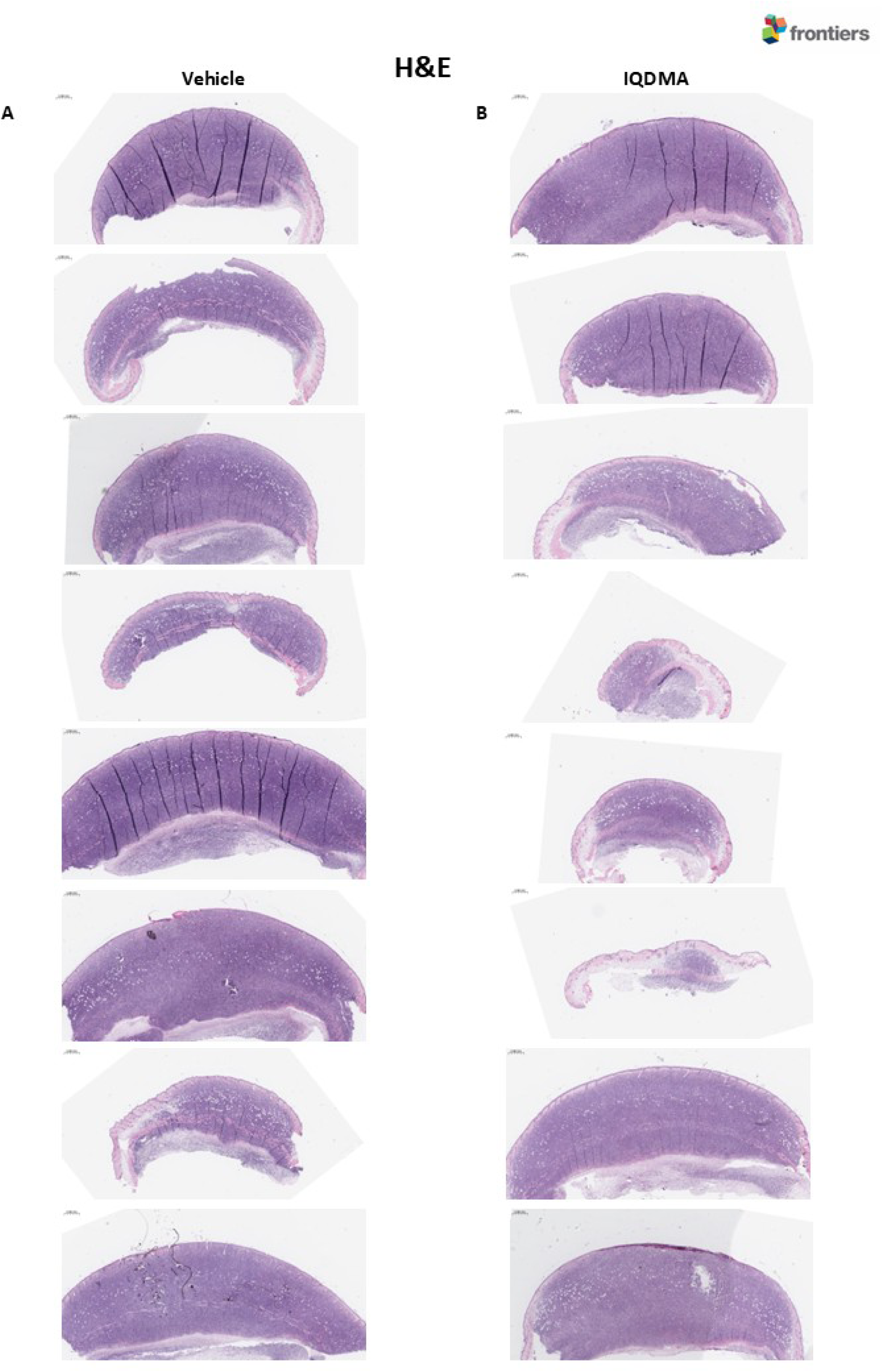
Histological examination of skin tumor tissues from a tumor-stage Mycosis Fungoides mouse model following IQDMA treatment. (A-B) Representative hematoxylin and eosin (H&E)-stained sections of skin tumor tissues from mice treated with vehicle **(A)** or 10 mg/kg IQDMA **(B)** administered daily via intraperitoneal injection

**Supplementary Figure 4:**
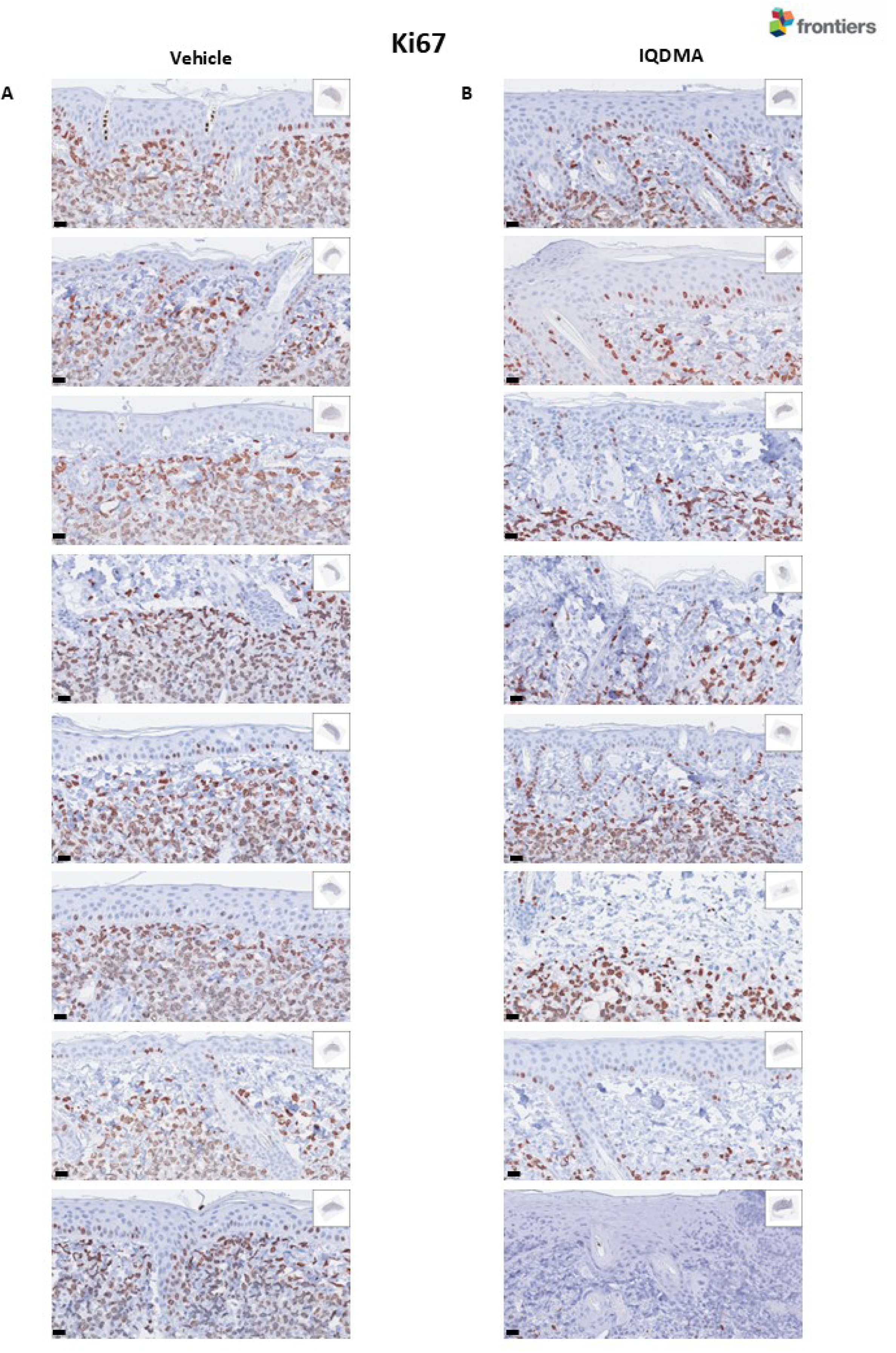
Immunohistochemical analysis of Ki-67 expression in skin tumor tissues. (A-B) Representative images of Ki-67 staining in skin tumor tissues from vehicle-treated **(A)** and IQDMA-treated **(B)** mice. Ki-67-positive cells are stained brown, while nuclei with no Ki-67 expression are counterstained blue—scale bars: 20 μm.

**Supplementary Figure 5:**
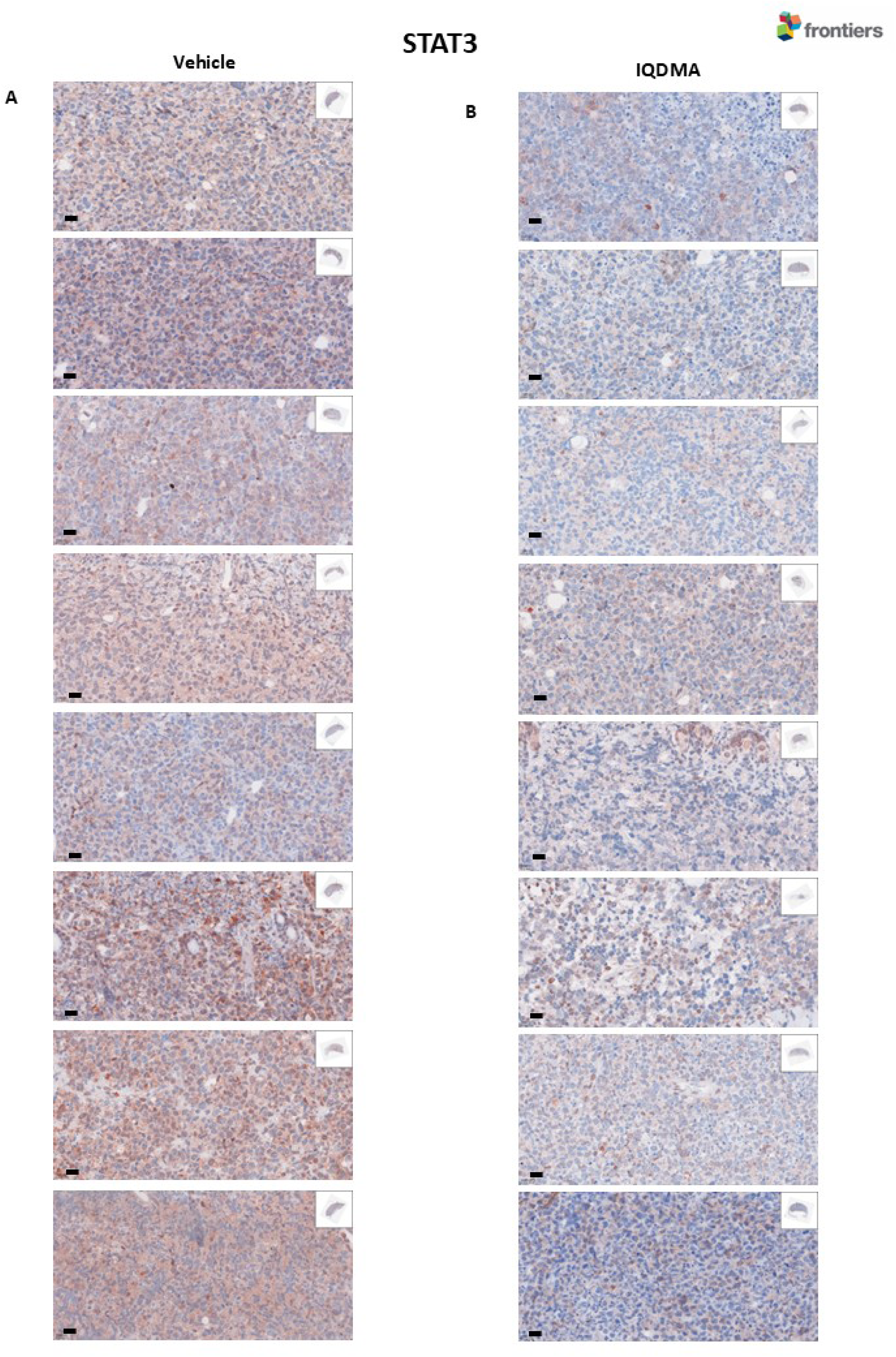
Immunohistochemical analysis of STAT3 expression in skin tumor tissues. (A-B) Representative images of STAT3 staining in skin tumor tissues from vehicle-treated **(A)** and IQDMA-treated **(B)** mice. STAT3-positive cells are stained brown. Scale bars: 20 μm.

**Supplementary Figure 6:**
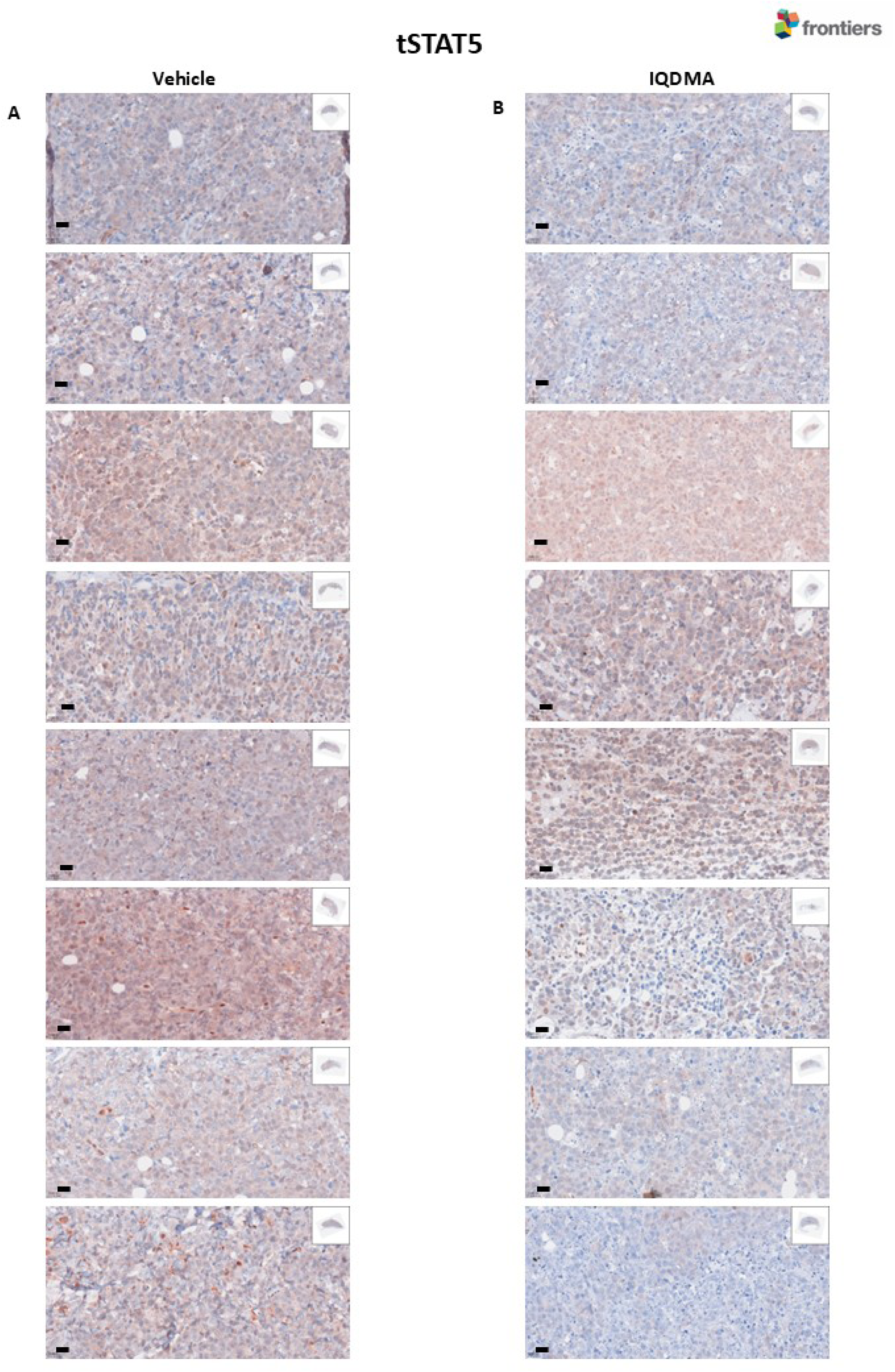
Immunohistochemical analysis of total STAT5 (tSTAT5) expression in skin tumor tissues. (A-B) Representative images of tSTAT5 staining in skin tumor tissues from vehicle-treated **(A)** and IQDMA-treated **(B)** mice. Scale bars: 20 μm.

**Supplementary Figure 7:**
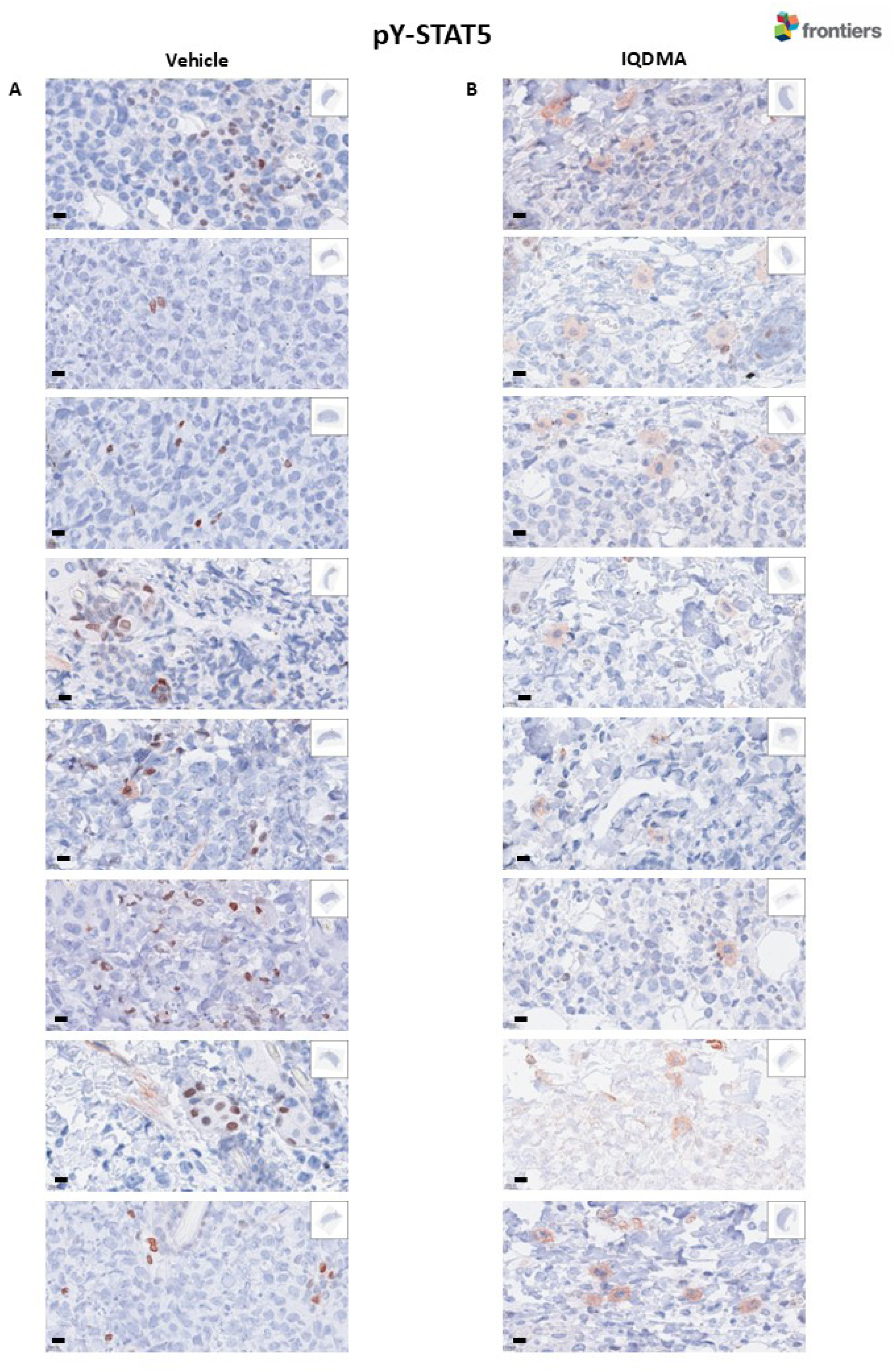
Immunohistochemical analysis of phosphorylated STAT5 (pY-STAT5) expression in skin tumor tissues**. (A-B)** Representative images of pY-STAT5 staining in skin tumor tissues from vehicle-treated **(A)** and IQDMA-treated **(B)** mice. pY-STAT5-positive cells are stained brown. A compartmental shift of pY-STAT5 from the nucleus to the cytoplasm was observed in the IQDMA-treated group—scale bars: 10 μm.

